# Mitotic exchange in female germline stem cells is the major source of *Sex Ratio* chromosome recombination in *Drosophila pseudoobscura*

**DOI:** 10.1101/2022.06.07.495109

**Authors:** Spencer Koury

**Author notes:** Address: 2613 Ashwood Avenue, Nashville, TN 37212.

## Abstract

*Sex Ratio* chromosomes in *Drosophila pseudoobscura* are selfish *X* chromosome variants associated with three non-overlapping inversions. In the male germline, *Sex Ratio* chromosomes distort segregation of *X* and *Y* chromosomes (99:1), thereby skewing progeny sex ratio. In the female germline, segregation of *Sex Ratio* chromosomes is mendelian (50:50), but non-overlapping inversions strongly suppress recombination establishing a 26 Megabase haplotype (constituting ~20% of the haploid genome). Rare crossover events located between non-overlapping inversions can disrupt this haplotype, and recombinants have sometimes been found in natural populations. We recently reported on the first lab-generated *Sex Ratio* recombinants occurring at a rate of 0.0012 crossovers per female meiosis. An improved experimental design presented here reveals these recombination events were 6.5-fold more frequent than previously estimated. Furthermore, recombination events were strongly clustered, indicating the majority arose from mitotic exchange in female germline stem cells and not from meiotic crossing-over in primary oocytes. Finally, recombination-induced viability defects consistent with unequal exchange caused asymmetric recovery rates of complementary recombinant classes. Incorporating these experimental results into population models for *Sex Ratio* chromosome evolution provided a substantially better fit to natural population frequencies and allowed maintenance of the highly differentiated 26 Megabase *Sex Ratio* haplotype without invoking strong epistatic selection. This study provides the first estimate of spontaneous mitotic exchange for naturally-occurring chromosomes in *Drosophila* female germline stem cells, reveals a much higher *Sex Ratio* chromosome recombination rate, and develops a mathematical model that accurately predicts the rarity of recombinant *Sex Ratio* chromosomes in natural populations.

## INTRODUCTION

Selfish sex chromosomes are a special class of segregation distorters found in organisms with chromosomal sex determination (Burt and Trivers 2006). Because non-mendelian transmission of the sex chromosomes skews the ratio of female to male offspring, these chromosomal variants are called *Sex Ratio* chromosomes. *Sex Ratio* (*X_SR_*) chromosomes are found in 16 species of Drosophila (an *XY* sex determination system) where they are typically associated with multiple inversions of the *X* chromosome (Jaenike 2001; Courret *et al.* 2019). In *Drosophila pseudoobscura*, all necessary and sufficient genes for the strong sex ratio distortion phenotype are linked to three non-overlapping inversions of the right arm of the metacentric *X* chromosome (Figure 1A) (Dobzhansky and Epling 1944; Wallace 1948).

**Figure 1.**
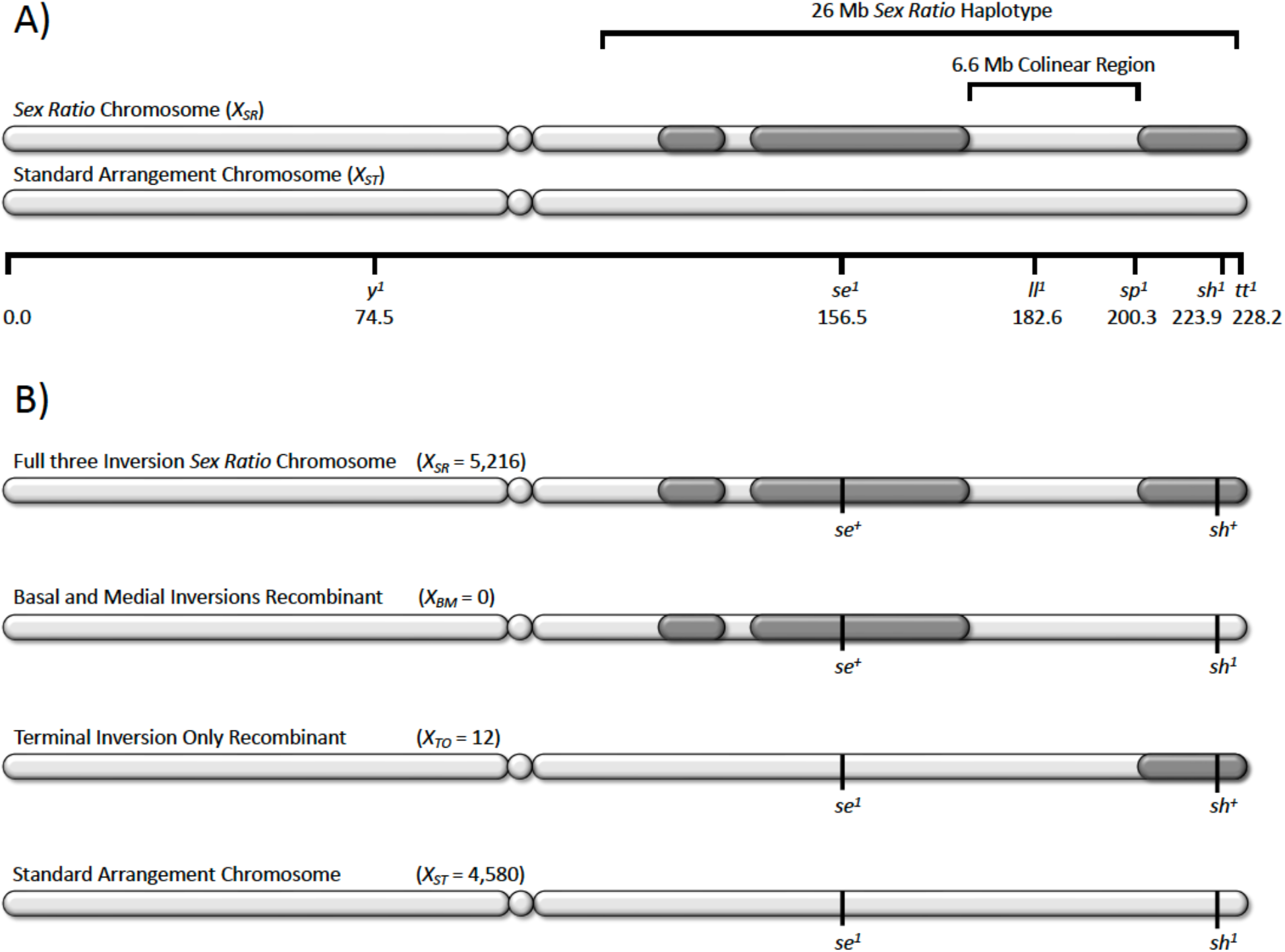
Schematic for recombination experiments with metacentric *X* chromosomes of *D. pseudoobscura.* A) Centromeres are depicted as centrally placed circles and the three non-overlapping inverted regions of the *Sex Ratio* chromosomes are shown in dark grey on the *X* chromosome right arm. Physical dimensions of *Sex Ratio* haplotype and colinear region are listed above the chromosome, below chromosomes are the genetic map position for the visible markers. B) Summary of Fuller *et al.* (2020) recombination experiment with raw counts provided in parentheses, the asymmetry of this result was very unexpected under a reciprocal-exchange model of meiotic crossing-over (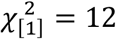, *p* = 2.85 × 10^-4^).

In natural populations of *D. pseudoobscura*, the three inversions of *X_SR_* chromosomes are found in near perfect linkage disequilibrium (*r*^2^ = 0.99) generating a single, large, highly differentiated 26 Mb *Sex Ratio* (*SR*) haplotype (Lewontin 1964; Fuller *et al.* 2020). Rare recombinant chromosomes that disrupt the three-inversion 26 Mb *SR* haplotype have been observed in natural population surveys (Wallace 1948; Beckenbach 1996). Cytogenetic analysis confirms these rare recombinants did not arise from ectopic exchange between *D. pseudoobscura X_SR_* and *Y* chromosomes; and more generally, brachycerous Dipterans have achiasmate male meiosis (Gethmann 1986; Beckenbach 1996). However, further detailed phenotypic study of recombinant *X_SR_* chromosomes has historically been complicated by their rarity and difficult maintenance in the lab (Wallace 1948).

We recently reported rare recombination (on the order of 1 in 1000 female meioses) occurring in the 6.6 Mb colinear region located between the medial and terminal inversions for heterozygotes of *Sex Ratio* and *Standard X* chromosomes (*X_SR_/X_ST_*) (Figure 1B) (Fuller *et al.* 2020). Segregation assays of recombinant chromosomes revealed the *X*-linked genetic architecture of the strong sex ratio distortion phenotype consists of at least two loci: a distorter gene in the proximal half of the right arm and a modifier gene in the distal half of the right arm. Using a simple model of gametic phase disequilibrium, we inferred strong epistatic selection must counteract recombination to maintain approximately 20% of the haploid genome as the single, highly differentiated *SR* haplotype observed in nature (Fuller *et al.* 2020). However, this conclusion was based on a model that did not account for the *X*-linked inheritance of *SR* haplotype, female-limited recombination or male-specific segregation distortion, and was highly sensitive to the experimentally determined recombination rate.

The recombination experiment reported in Fuller *et al.* (2020) followed standard testcross protocol mass crossing 20 female *X_SR_/X_ST_* heterozygotes to 20 tester strain males in 6 fl. oz milk bottles (Bridges and Brehme 1944). For logistical reasons, only male progeny were scored for recombination of *X*-linked visible markers before being discarded. Twelve of the nearly ten thousand males scored were recombinants; however, all 12 males exhibited the same marker combination (indicating a recombinant carrying only the terminal inversion of *X_SR_* chromosomes, denoted *X_TO_*) and we observed none of the complementary recombinant *X_SR_* chromosome (carrying both the basal and medial inversions but without the terminal inversion, denoted *X_BM_*). Under a reciprocal-exchange model of meiotic crossing-over, this was a very rare and unexpected result (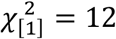, *p* = 2.85 × 10^−4^). Further screening (>10,000 individuals) demonstrated stable *X_BM_* stocks could indeed be established, and polytene chromosome squashes confirmed the recombination events occurred in the 6.6 Mb colinear region located between the medial and terminal inversions of the *SR* haplotype (see photomicrograph in Supplemental Figure 7 of Fuller *et al.* 2020).

In the Fuller *et al.* (2020) experimental design, the *short^1^* mutation marked the absence of the terminal inversion of *X_SR_* chromosomes. Only male progeny were scored for recombination due to incomplete penetrance of the wing vein mutation *short*^1^ in female homozygotes. Any residual penetrance problems in hemizygous males could have caused incorrect classification of the missing recombinant *X_BM_* chromosomes as non-recombinant wildtype *X_ST_* chromosomes. Furthermore, if the missing recombinant *X_BM_* chromosomes have strong recessive viability defects, then these recombinants would not have been detected in hemizygous male progeny. Finally, in a ten thousand fly experiment, other rare events (*e.g.*, gene conversion or somatic mutation) could have inflated the estimated crossover rate. To address these potential sources of bias, a new recombination experiment was conducted using single-female crosses, scoring both male and female progeny, and retaining recombinant progeny for confirmation testcrosses.

Beyond the potential design artifacts listed above, the rare *X_SR_/X_ST_* recombinants may not have been the product of meiotic crossing-over, but rather arose from mitotic exchange in the female germline stem cells (GSC) prior to meiosis. This biological source of variation would confound normal meiotic crossover rate estimation with a recombination process that precedes meiosis. In Drosophila, ovaries are divided into 16-20 ovarioles, each containing 2-3 GSCs occupying a niche in the anterior region of the ovariole (*i.e.*, the germarium, as shown in Figure 2). Self-renewing GSCs divide asymmetrically to produce a differentiating cystoblast which undergoes four rounds of mitosis with incomplete cytokinesis to form a 16-cell cyst (reviewed in McLaughlin and Bratu 2015). In a well-studied determination process, only one of these sixteen cells will develop into an oocyte and undergo meiosis while the other fifteen cells become nurse cells that do not transmit genetic material to the next generation (reviewed in Huynh and St Johnston 2004). Therefore, recombinant progeny resulting from canonical meiotic crossing-over (reciprocal-exchange) in each oocyte are causally independent of meiotic crossing-over in other oocytes. In contrast, mitotic exchange occurring in self-renewing stem cells has the potential to create clusters of non-independent (clonal) recombinant progeny, all originating from a single GSC mitotic event (Figure 2).

**Figure 2.**
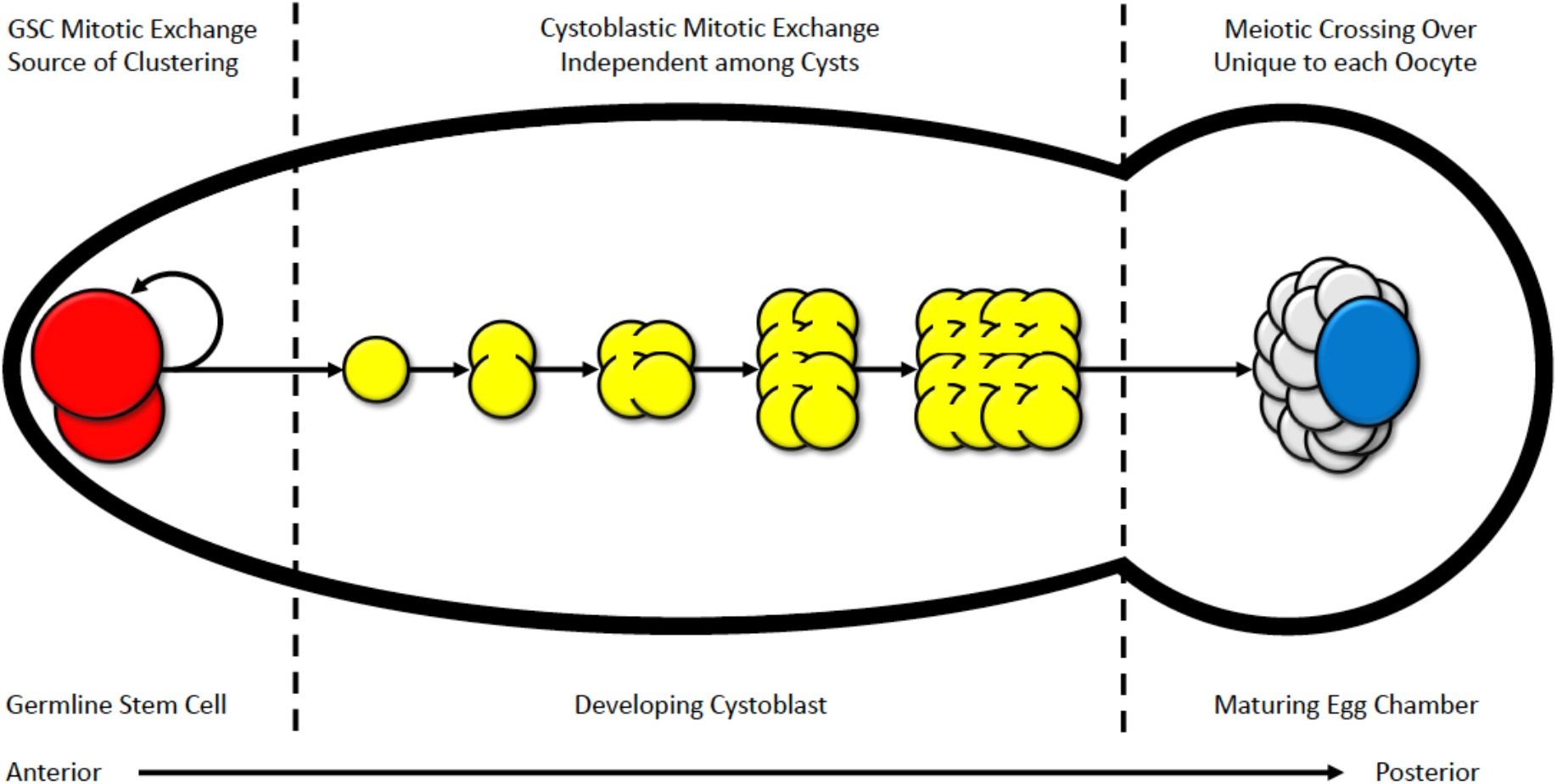
Cartoon sequence of germarium and single egg chamber development in anterior region of Drosophila ovariole. Mitotic exchange in germline stem cells (GSC, red) with their asymmetric division and self-renewal (reflexive arrow) can cause clustering in recombination datasets. Mitotic exchange in developing cystoblast (yellow) cannot cause clustering as only one of sixteen nuclei will develop into the oocyte (blue), with the other 15 becoming nurse cells (grey). Meiotic crossing-over in oocyte (blue) only occurs after development of the 16-cell cyst (yellow) and are causally independent among oocytes. Therefore, clustering of recombination events must originate from mitotic exchange in GSC or earlier in development. Cartoon does not represent a proportional scale but is meant to illustrate the sequential events allowing for interpretation of clustering in recombination data.

Mitotic exchange in the female germline cells is difficult to measure because crossing-over in prophase of meiosis I masks this signal under normal conditions (Ashburner *et al.* 2004). Nevertheless, 18% of *X-Y* exchanges from compound chromosome *C(1)RM/Y* in female meiosis were attributed to GSC mitotic exchange (Neuhaus 1936), and rates of female GSC mitotic exchange have been elevated by X-ray irradiation (Bateman and Chandley 1965). In *Drosophila pseudoobscura X_SR_/X_ST_* heterozygotes, the non-overlapping inversions of *X_SR_* chromosomes strongly suppress meiotic crossing-over on the entire right arm of the *X* chromosome; thereby presenting a unique opportunity to measure spontaneous mitotic exchange in female GSCs for naturally-occurring *X* chromosomes under normal conditions. In this specific context, GSC mitotic exchange can be detected statistically by the clustered distribution (*i.e.*, over-dispersion) of recombination events.

In the present study, an improved experimental design revealed recombination events were substantially more frequent than previously estimated. Statistical analysis rejected the hypothesis that meiotic crossing-over generated the present recombination dataset. Three distinct types of clustering in recombination events indicated that the non-independence in this dataset arose from unequal mitotic exchange in the female GSCs. To understand the implications of these experimental results for natural populations, the Edwards (1961) population genetic model of *Sex Ratio* was extended to incorporate recombination in an Appendix to this paper. Modeling unequal mitotic exchange in the *Sex Ratio* system provided a substantially better fit to natural population frequencies of *X_SR_* chromosomes in *D. pseudoobscura* and, despite the higher recombination rate, the model was able to explain the relative rarity of recombinant *X_BM_* and *X_TO_* chromosomes in the wild.

## MATERIALS AND METHODS

### Live Stock Construction

The three focal *Sex Ratio* (*X_SR_*) chromosomes were identified from natural population collections in Zion National Park, UT, USA in September 2013 and Kaibab National Forest, AZ, USA in September 2017. Subsequently in 2018, the base *X_SR_* stocks were created by re-isolation of *X_SR_* chromosomes and twenty generations of backcrossing to three different highly inbred stocks (*F* > 0.99), each with a multiply-marked standard arrangement *X* chromosome (*X_ST_*). The resulting nine stocks (three *X_SR_* chromosome isolates on each of three different backgrounds) were segregating for an unmarked *X_SR_* and a marked *X_ST_* on a homozygous genetic background.

The visible markers of the *X_ST_* chromosomes spanned the whole right arm of the metacentric *X* chromosome; and in combination, the mutations *yellow^1^* (*y^1^*, 1-74.5), *sepia^1^* (*se^1^*, 1-156.5), *lanceolate^1^* (*ll^1^*, 1-182.6), *snapt^1^* (*sp^1^*, 1-200.3), *short^1^* (*sh^1^*, 1-225.9), and *tilt^1^* (*tt^1^*, 1-228.2) allowed detection of rare *X_SR_* recombinants events (Orr 1995; Fuller *et al.* 2020). Specifically, visible mutation combinations *se^1^ sh^1^*, and independently *se^1^ sp^1^ tt^1^*, were introgressed onto the *D. pseudoobscura* reference genome background (*F* > 0.99). These two independent tester strains were used for outcrossing procedures in experiments to identify recombination events. A full list of *X_SR_* stocks, *X_ST_* inbred lines, the visible mutations they carry, and the National Drosophila Species Stock Center identification for their genetic background is provided in Table 1 and as a Reagent Table in File S1.

**Table 1.** A complete list of lines used. All lines have highly inbred genetic backgrounds with National Drosophila Species Stock Center identifier listed. Inbred lines and tester strains are homozygous for standard arrangement *X* chromosomes. *Sex Ratio* stocks are segregating for the inverted arrangement unmarked *X_SR_* and standard arrangement multiply-marked *X_ST_*.

| ***Sex Ratio* ($X_{SR}$) Segregating Stocks** | **Visible Markers** | **Genetic Background** |
| --- | --- | --- |
| *Sex Ratio* Isolate KBNP2 | $y^1, se^1, sh^1$ | 14011-0121.06 |
| *Sex Ratio* Isolate KBNP2 | $se^1, ll^1, sp^1, tt^1$ | 14011-0121.08 |
| *Sex Ratio* Isolate KBNP2 | $se^1, sh^1$ | Phadnis Lab Line 2020 |
| *Sex Ratio* Isolate Z8 | $y^1, se^1, sh^1$ | 14011-0121.06 |
| *Sex Ratio* Isolate Z8 | $se^1, ll^1, sp^1, tt^1$ | 14011-0121.08 |
| *Sex Ratio* Isolate Z8 | $se^1, sh^1$ | Phadnis Lab Line 2020 |
| *Sex Ratio* Isolate Z6 | $y^1, se^1, sh^1$ | 14011-0121.06 |
| *Sex Ratio* Isolate Z6 | $se^1, ll^1, sp^1, tt^1$ | 14011-0121.08 |
| *Sex Ratio* Isolate Z6 | $se^1, sh^1$ | Phadnis Lab Line 2020 |
| **Standard Arrangement ($X_{ST}$) Inbred Lines** | **Visible Markers** | **Genetic Background** |
| Standard Inbred Line 1 | $y^1, se^1, sh^1$ | 14011-0121.06 |
| Standard Inbred Line 2 | $se^1, ll^1, sp^1, tt^1$ | 14011-0121.08 |
| Standard Inbred Line 3 | $se^1, sh^1$ | Phadnis Lab Line 2020 |
| **Standard Arrangement ($X_{ST}$) Tester Strains** | **Visible Markers** | **Genetic Background** |
| Standard Tester Strain 1 | $se^1, sh^1$ | 14011-0121.94 |
| Standard Tester Strain 2 | $se^1, sp^1, tt^1$ | 14011-0121.94 |

### Recombination Experimental Design

To test for recombination in the 6.6 Mb colinear region located between medial and terminal inversions of *D. pseudoobscura X_SR_* chromosomes, a single-block, fully-coded, randomized design was conducted with the experimenter blinded to genotypic treatment. The factorial design incorporated three independent *X_SR_* chromosome isolates, each on three independent autosomal genetic backgrounds, and three independent multiply-marked *X_ST_* chromosomes on those same three autosomal genetic backgrounds (3 × 3 × 3). Experimental *F_1_* female heterozygotes (*X_SR_/X_ST_*) were generated by performing all pairwise crosses, excluding genotypes that would produce inbred genetic backgrounds (3 × 3 × 3 – 9), for a total of 18 unique outbred experimental genotypes (Table S1 in File S2).

A total of 180 *F_1_* experimental virgin females and 900 tester strain males were collected within 12 hours of eclosion over a three-day period. Females and males were aged in separate vials for an additional three-day period. Using light CO2 anesthesia, ten females for each genotypic treatment were then individually mated to five tester males possessing an independent genetic background (from the *D. pseudoobscura* reference genome) and the respective *X*-linked markers (*se^1^ sh^1^* or alternately *se^1^ sp^1^ tt^1^).* The 180 single-female crosses (18 genotypic treatments with 10 replicates each) were allowed another three-day CO2 recovery period, during which all experimental crosses were coded and randomized. Experimental crosses were maintained at 21°C, 65% relative humidity, and 14:10 hour light:dark cycle. The single-female crosses were individually tap transferred onto experimental food for a three-day egglaying period that occurred when all flies were minimally one-week and no older than two-weeks post-eclosion. Experimental food was standard cornmeal molasses Drosophila media seeded with live yeast. After three days, the parental adult were removed under CO2 anesthesia and the food was hydrated as needed with 0.5% v/v propionic acid.

Both male and female *F_2_* progeny were scored daily for visible mutations (*se^1^*, *sh^1^* or alternately *se^1^*, *sp^1^*, *tt^1^*) until adult flies stopped emerging, for a total of 17,604 progeny. However, scoring recombination between *se^1^* and *sp^1^ tt^1^* in female progeny was confounded by the presence of *ll^+^* in the tester strain’s colinear region that altered penetrance of wing vein mutations *sp^1^* and *tt^1^*. Therefore, the six genotypic treatments that used the *se^1^*, *sp^1^, tt^1^* marker system could not be reliably scored and were excluded from analysis.

When a putative *F_2_* recombinant was identified, this individual was isolated and mated to multiply-marked individuals (*se^1^*, *sh^1^*) of the opposite sex to establish a temporary stock. Confirmation that the individual carried a true recombinant *X* chromosome was provided by transmission and detection of that same marker combination in the subsequent generation. Initial *F_2_* screening identified 102 recombinants among 11,808 progeny scored for visible markers *se^1^* and *sh^1^*. Confirmation tests of these 102 recombinants revealed 10% resulted from errors in scoring the wing vein marker *sh^1^*. Rates of misclassification for *sh^1^* as wildtype *sh^+^* (3/24, exact binomial 95% confidence interval 0.03-0.36) were statistically indistinguishable from the rate of misclassifying *sh^+^* as a mutant *sh^1^* (7/78, exact binomial 95% confidence interval 0.03-0.18).

### Statistical Analysis of Recombination

An ANOVA with type II sum of squares was conducted testing *X_SR_* chromosome isolate effect and genetic background effect on the observed recombination rate. First, any putative recombinant that was not confirmed in subsequent generations was recoded in the dataset. Second, the recombination rate was calculated as the number of confirmed recombinants divided by the total number of progeny scored for each single-female cross. Third, this proportion was arcsine-transformed 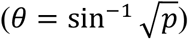 to fit ANOVA’s error term normality assumption (Sokal and Rohlf 1995). The linear model analyzed was *θ_ijk_* = *γ_i_* + *τ_j_* + *ε_ijk_*; where *γ_i_* is the effect of the *i^th^ X_SR_* chromosome isolate, *τ_j_* is the effect of the *j^th^* genetic background, and *ε_ijk_* represents the error term. Because no statistically significant effects related to *X_SR_* chromosome isolate or genetic background were detected, all subsequent analyses did not subdivide the dataset based on these factors.

To investigate whether the observed data were consistent with meiotic crossing-over, the count of recombinants per single-female cross was compared to the Poisson distribution. Assuming meiotic recombination events were rare and occur independently, then the probability of observing a given number of recombinants (*x_i_*) for single-female cross *i* with *n_i_* offspring is 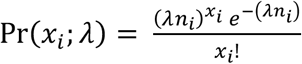, where *λ* is the experiment-wide rate of recombination. The probability distributions for each single-female cross were combined to generate an experiment-wide distribution accounting for differences in offspring number among crosses. To evaluate the full observed dataset, *χ*^2^ goodness-of-fit test with four degrees of freedom was conducted with expected values given by the experiment-wide distribution. The probability of observing a single-female cross with more than four recombinants in this design is always less than 0.01. Therefore, the few crosses that produced 5, 6, and 9 recombinants were pooled into a single class consisting of ≥ 4 recombinants (this isa statistically conservative procedure for this dataset).

To further investigate whether the observed data were consistent with reciprocal-exchange model of meiotic crossing-over, the proportion of recombinants per single-female cross carrying only the terminal inversion (*X_TO_*), and not the basal or medial inversions (*X_BM_*), was compared to the binomial distribution. Assuming recombination events were independent with an equal probability of transmitting either complementary recombinant *X* chromosome to progeny (*p* = (1 – *q*) = 0.5), then the probability of observing a given number of recombinants *k_i_* for single-female cross *i* carrying only the terminal inversion is binomially distributed 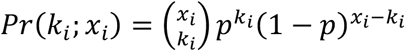 where *x_i_* is the number of recombinants from cross *i* with *k_i_* total progeny scored. The probability distribution for each single-female cross producing two or more recombinants was binned to generate an experiment-wide distribution that controlled for differences in recombinant numbers among crosses. The observed data were then compared to the binned expected distribution with a *χ^2^* goodness-of-fit test with four degrees of freedom.

Finally, to investigate whether sex of the progeny scored for recombination had an effect on the chances of observing complementary recombinant *X* chromosomes, a 2 × 2 contingency table was analyzed. Under a reciprocal-exchange model of meiotic crossing-over with no effect of progeny sex on detection probability, all four classes (*i.e.*, females with *X_BM_*, males with *X_BM_*, females with *X_TO_*, and males with *X_TO_*) were expected to occur with equal frequency. The observed data were compared to this expectation (discrete uniform distribution) using a *χ^2^* goodness-of-fit test with three degrees of freedom.

### Data Availability

*D. pseudoobscura* stocks listed in Table 1 and further described in Reagents Table in File S1 are available upon request. The author affirms that all data necessary for confirming the conclusions of the article are present within the article, figures, and tables. Raw counts from the recombination experiments are provided as a supplement in File S3.

## RESULTS

After adjusting for false positives, the observed experiment-wide recombination rate was 0.00779 (92 recombinants in 11,808 progeny). Substantial heterogeneity among the single-female crosses was observed in recombination rates (for raw counts see File S3). The dataset was generated under common garden conditions as a single-block, fully-randomized, full-factorial experimental design with the investigator blinded to treatment, such that the observed heterogeneity was not likely an artifact of design. To investigate the biological sources of recombination rate heterogeneity, and to test the null hypothesis that a reciprocal-exchange model of meiotic crossing-over produced this pattern of heterogeneity, the distribution of observed data was subjected to further statistical analysis.

### No Genetic Variation for Recombination Rate

To investigate whether the *X_SR_* chromosome isolate or genetic background affected recombination, a type II ANOVA was conducted on arcsine transformed rate data. No statistically significant effects were detected for the three *X_SR_* chromosome isolates (*F*_2,115_ = 0.57, *p* = 0.57) or the three genetic backgrounds (*F*_2,115_ = 0.84, *p* = 0.44) (Table 2). Additionally, no statistically significant effects were detected in sex-stratified analysis of the rate data (Table S2 in File S2). Therefore, all subsequent analyses did not subdivide dataset based on *X_SR_* chromosome isolate or genetic background.

**Table 2.** Recombination rate ANOVA table. No genetic variation for recombination rate was detected when analyzing the effect of three *X_SR_* chromosome isolates and three heterozygous genetic backgrounds with type II ANOVA.

| Source of Variation | *df* | *SS* | *MS* | *F<sub>s</sub>* | *p*-value |
| --- | --- | --- | --- | --- | --- |
| $X_{SR}$ Chromosome Isolate | 2 | 19.95 | 9.98 | 0.57 | 0.57 |
| Genetic Background | 2 | 29.16 | 14.58 | 0.84 | 0.44 |
| Residuals | 115 | 2005.82 | 17.44 |  |  |
| Total | 119 | 2054.93 |  |  |  |

### Recombination Events were Clustered Among Females

To investigate whether the data were consistent with meiotic crossing-over (*i.e.*, independence of recombination events among single-female crosses), the observed count of recombinants per single-female cross was compared to the Poisson distribution using the experiment-wide recombinant frequency as the rate parameter λ (Figure 3A). A *χ*^2^ goodness-of-fit test with four degrees of freedom revealed that the observed recombination events were not independent (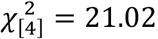, *p* = 1.43 × 10^-4^) (Table 3). Substantially more single-female crosses produced either no recombinants or multiple recombinants than were expected assuming independence of recombination events. The coefficient of dispersion 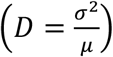 differed with statistical significance from the Poisson distribution’s expected value of one (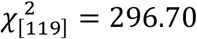, *p* = 1.09 × 10^-17^), indicating the data were strongly clustered. The same pattern of over-dispersion (clustering) was present in sex-stratified analysis of observed data (Table S3 in File S2). Strong clustering of observed data rejects the null hypothesis that reciprocal-exchange in meiosis produced this dataset and suggests that multiple recombinant *X* chromosomes recovered from the same experimental *F_1_* single-female cross were clonal.

**Figure 3.**
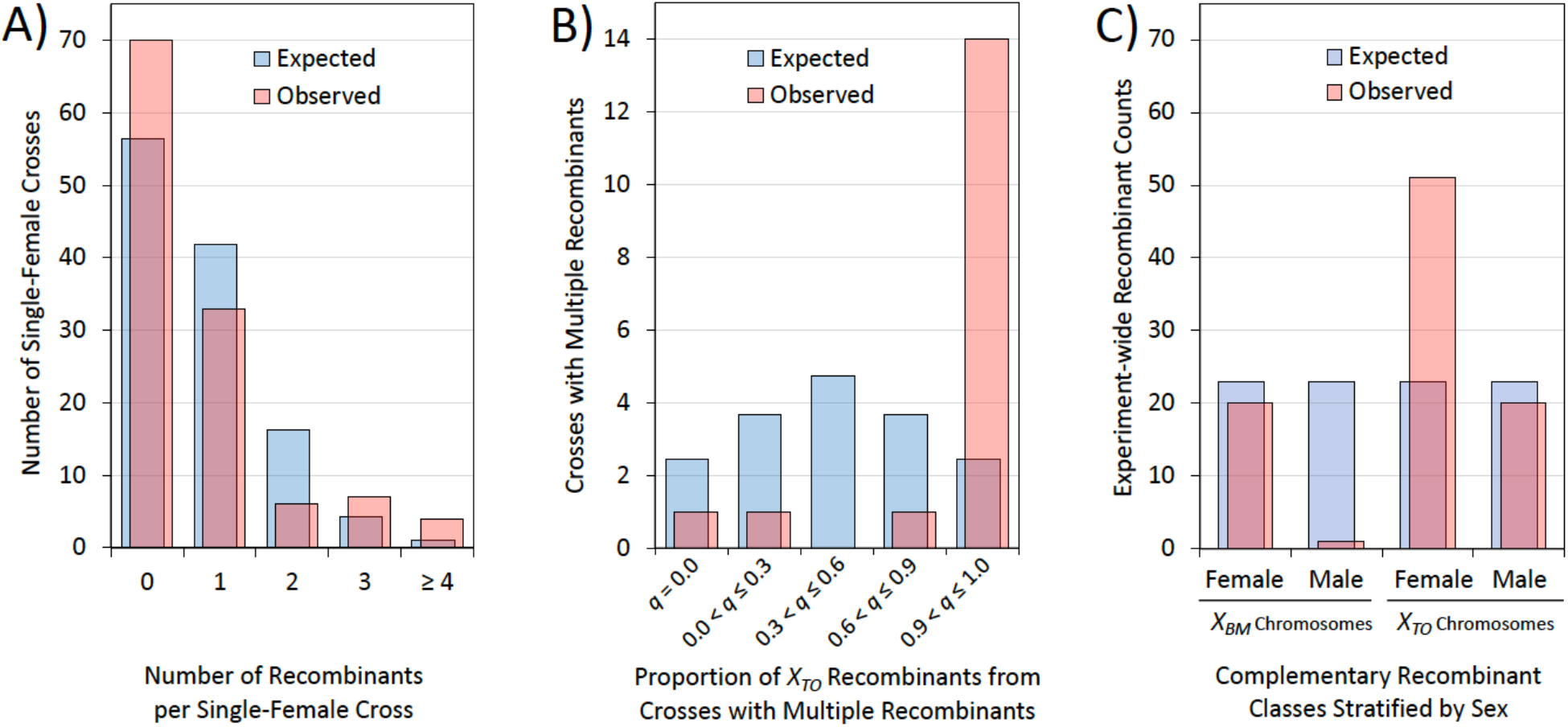
Observed versus expected distribution of recombination events. A) Observed recombination events (red) were clustered among single-female crosses when compared to Poisson expectations (blue) based on experiment-wide recombination rate (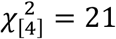, *p* = 1.43 × 10^-4^) (see Table 3). B) Proportion of recombinants (*q*) carrying *X_TO_* were clustered within single-female crosses when compared to expected proportions from the binomial distribution. Multiple recombinants from single-female crosses (red) were more likely to all be *X_TO_* than expected by chance (blue) 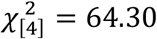, *p* = 1.75 × 10^-13^) (see Table 4). C) Recovery rates of complementary recombinants were neither equivalent between nor independent of *F_2_* progeny sex. The observed data (red) differed from uniform discrete expectations (blue) with statistical significance (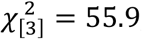, *p* = 2.15 × 10^-12^) (see Table 5). A deficit of both complementary recombinants (*X_TO_* and *X_BM_*) was detected in the hemizygous (male) state and was particularly severe for *X_BM_*.

**Table 3.** Recombination events were not independent among *F_1_* single-female crosses. The number of recombination events per single-female cross, if events were independent, should be Poisson distributed. A *χ*^2^ goodness-of-fit test revealed the observed recombination events per single-female cross differed with statistically significance from Poisson expectations using a experiment-wide recombination rate (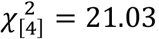, *p* = 1.43 × 10^-4^). Clustering was even more extreme than the *χ*^2^ test indicates due to conservative procedure of binning crosses producing four or more recombinants (see Figure 3A).

| Category | *Expected* | *Observed* | $(Obs-Exp)^2/Exp$ |
| --- | --- | --- | --- |
| 0 Recombinants | 56.44 | 70 | 3.26 |
| 1 Recombinant | 41.87 | 33 | 1.88 |
| 2 Recombinants | 16.24 | 6 | 6.46 |
| 3 Recombinants | 4.36 | 7 | 1.60 |
| $\geq 4$ Recombinants | 1.09 | 4 | 7.83 |
| | | | $\chi^2 = 21.03$ |

**Table 4.** Multiple recombination events were not independent within *F_1_* single-female crosses. Proportion of recombinants (*q*) carrying one inversion complement (*X_TO_*) in single-female crosses producing multiple recombinants *via* meiotic crossover should be binomially distributed. A *χ*^2^ goodness-of-fit test revealed statistically significant deviation from these expectations (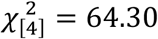, *p* = 1.75 × 10^-13^). Multiple recombinants from single-female crosses (predominantly *X_TO_*) were much more likely to carry the same inversion complement than expected by chance (see Figure 3B).

| Category | Expected | Observed | (Obs-Exp)<sup>2</sup>/Exp |
| --- | --- | --- | --- |
| $q = 0.0$ | 2.44 | 1 | 0.85 |
| $0.0 < q \leq 0.3$ | 3.69 | 1 | 1.96 |
| $0.3 < q \leq 0.6$ | 4.74 | 0 | 4.74 |
| $0.6 < q \leq 0.9$ | 3.69 | 1 | 1.96 |
| $q = 1.0$ | 2.44 | 14 | 54.79 |
| | | | $\chi^2 = 64.30$ |

**Table 5.** Recovery rates of complementary recombinants were not equivalent among *F_2_* progeny sex: The observed recombinants classified by sex and inversion complement differed from uniform discrete expectations with statistical significance (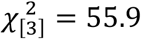, *p* = 2.15 × 10^-12^) (see Figure 3C). Observation of complementary recombinants was strongly dependent on the *F_2_* sex in which they were detected, with the most noticeable deficit occurring in the male *X_BM_* class.

| Category | Expected | Observed | (Obs-Exp)<sup>2</sup>/Exp |
| --- | --- | --- | --- |
| $X_{BM}$, Female | 23 | 20 | 0.39 |
| $X_{BM}$, Male | 23 | 1 | 21.04 |
| $X_{TO}$, Female | 23 | 51 | 34.09 |
| $X_{TO}$, Male | 23 | 20 | 0.39 |
| | | | $\chi^2 = 55.91$ |

### Multiple Recombination Events were Clustered Within Females

To investigate whether the observed data were consistent with meiotic crossing-over (*i.e.*, independence of recombination events within a single-female cross), the proportion of recombinants (*q*) from a single-female cross that carried *X_TO_* was compared to the binomial distribution (Figure 3B). A *χ*^2^ goodness-of-fit test revealed multiple recombinants from a single-female cross differed with statistical significance from binned expectations assuming independence (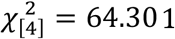, *p* = 1.75 × 10^-13^) (Table 4). The same pattern was present in sex-stratified analysis of data (Table S4 in File S2). Multiple recombinants from a given single-female cross were more likely to carry the same inversion complement than expected by chance, again rejecting the null hypothesis of meiotic reciprocal-exchange and further suggesting that multiple recombinant *X* chromosomes recovered in from the same experimental *F_1_* single-female cross were clonal.

### Recovery Rates of Recombinants were Neither Equivalent nor Independent of Sex

To investigate whether the sex of progeny scored for recombination had an effect on the probability of observing a recombinant (*i.e.*, recombination events in *F_1_* female germline were independent of *F_2_* progeny sex), the observed data were compared to a discrete uniform distribution (Figure 3C). The observed data differed from expectations with statistical significance (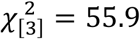, *p* = 2.15 × 10^-12^) (Table 5). These results demonstrated that recovery of recombinant *X* chromosomes was strongly dependent on the *F_2_* sex in which they were detected, with the most noticeable deficit occurring in the male *X_BM_* class.

The observed recovery rates for *X_BM_* recombinants in both males and females were much lower when compared to the complementary recombinant *X_TO_*. *χ*^2^ tests of independence failed even when different rates of production were estimated from the data for complementary recombinant classes (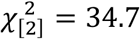, *p* = 1.06 × 10^-7^) (Table S5 in File S2). There was also a deficit of both complementary recombinants (*X_TO_* and *X_BM_*) detected in the hemizygous (male) state compared to the protected heterozygous (female) state. Again, *χ*^2^ tests of independence failed after different rates for both sexes and complementary classes were estimated from the data (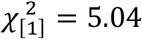, *p* = 0.0143) (Table S5 in File S2). Even after these corrections, the most noticeable deficit was still due to the hemizygous (male) *X_BM_* class, a genotype with notably reduced viability in the laboratory (Wallace 1948; Fuller *et al.* 2020; unpublished results). The process of recombination itself causing asymmetric recessive viability defects is consistent with a model of mitotic exchange generating a chromosomal deletion for *X_BM_*, and a complementary duplication for *X_TO_*, *via* unequal crossing-over in the female germline stem cell.

## DISCUSSION

The present results demonstrate that recombination events located between non-overlapping inversions of *Sex Ratio* chromosome heterozygotes in *D. pseudoobscura* were not as rare as previously estimated. Importantly, the recombination events exhibited strong clustering both among and within single-female crosses. The hypothesis that canonical meiotic crossing-over (reciprocal exchange) generated this recombination dataset was unambiguously rejected (Figure 3). At least 17 of 50 single-female crosses that produced recombinants can be statistically attributed to mitotic exchange in female germline stem cells (GSC) or at earlier stage in development (Figure 2). As a result, at least 64% of the total recombinant progeny observed in this experiment (59/92) originated from female germline mitotic exchange. The remaining 36% of recombinants occurred in single-female crosses that produced only one recombinant in the three-day egglaying period, and therefore provided insufficient information to determine whether their origin was due to mitotic exchange or meiotic crossing-over.

The observed experiment-wide recombination rate of 92 recombinants per 11,808 progeny scored (0.00779) was within the expected range if a single *X_SR_/X_ST_* heterozygous GSC was converted *via* mitotic exchange (expected range 0.00417-0.00781, calculated assuming 2-3 GSCs per ovariole, 16-20 ovarioles per ovary, 2 ovaries per female). In principle, mitotic exchange may occur after asymmetric division of GSCs in the cystoblast; but because each cyst only yields one oocyte, any mitotic exchanges subsequent to differentiation from GSCs cannot be the source of clustering (Figure 2). Conversely, the most extreme single-female recombination rates observed in this experiment (*e.g.*, 6/47 and 9/122) may be caused by mitotic exchange in primordial germ cells prior to their development into GSCs and therefore leading to a subpopulation of clonal GSCs with the converted genotype. Based on the close match with the expected range, the large majority of the observed recombination events appear to have occurred as mitotic exchange in female GSCs rather than in an earlier stage.

The statistical approach of this study cannot provide direct evidence of unequal mitotic exchange, but asymmetric patterns in the dataset (*i.e.*, specific deficit of *X_BM_* recombinants or deficit of hemizygous recombinants more generally) were consistent with unequal exchange. The most parsimonious explanation for the different recovery rates of complementary recombinants, and the observed recombination-induced recessive viability defects, is unequal exchange generating a chromosomal deletion for *X_BM_* and a complementary duplication for *X_TO_*. However, it is possible more complex chromosomal aberrations stemming from the mitotic exchange may have caused the disproportionate deficit of hemizygous *X_BM_* recombinants observed in Figure 3C. In future studies, identification of recombination breakpoints with long-read sequencing could provide direct evidence for this scenario’s predictions that: 1) clustered recombinants had clonal origins, 2) repetitive genetic elements mediated ectopic exchange, and 3) deleted and/or duplicated regions at breakpoints caused the observed fitness defects.

### Modeling Recombinant Rarity in Natural Populations

The higher recombination rate observed in this study intuitively suggests recombinant *Sex Ratio* chromosomes should be more frequent in the wild (greater than 10% of *X* chromosome variants). However, both *X_TO_* and *X_BM_* are still quite rare (less than 1%) in natural population samples (Sturtevant and Dobzhansky 1936; Wallace 1948; Beckenbach 1996). To investigate whether the properties of unequal mitotic exchange could explain the low frequencies of *X_TO_* and *X_BM_* in the wild, it was necessary to introduce recombination into the Edwards (1961) population genetic model of *Sex Ratio* in *D. pseudoobscura* (see Appendix for model extension).

Briefly, Edwards (1961) developed a sex-specific, genotypic model for *Sex Ratio* because *X*-linked meiotic drivers cause sustained difference in allele frequencies between the sexes. The extended model uses a mating table as a transition matrix to convert the 40 possible mating types (4 male genotypes × 10 female genotypes) to the next generation’s 14 genotypic frequencies (4 male genotypes and 10 female genotypes). The full 40 × 14 mating table is provided as File S4 and is rewritten in the Appendix as a system of recursion equations with 14 variables representing the frequency of each genotype. The genotypic frequencies in the next generation are a function of the frequencies in the previous generation and parameters *v_i,j_* for relative viability, *f_i,j_* for relative fertility, *k_i_* for transmission in the male germline, and *c_i,j_* for the recombination rate of double heterozygotes in the female germline (Equation A1 in Appendix).

I parameterized the extended model of Edwards (1961) with experimental data on viability, fertility, distortion, and recombination (Curtsinger and Feldman 1980; Fuller *et al.* 2020; unpublished results; this study). Making the additional assumption that fitness effects in recombinants are both recessive and proportional to the fraction of 26 Mb *Sex Ratio* haplotype retained after recombination, experiment-based estimates for all 34 parameters in this extended population genetic model were provided (Table A1 in Appendix). The equilibrium frequencies of all 14 genotypic variables were then solved for by iterating the system of equations. A range of initial frequency combinations between 0.01 and 0.99 were considered and all converged on values reported in Table 6 (column “Meiotic Crossing-Over”).

**Table 6.**
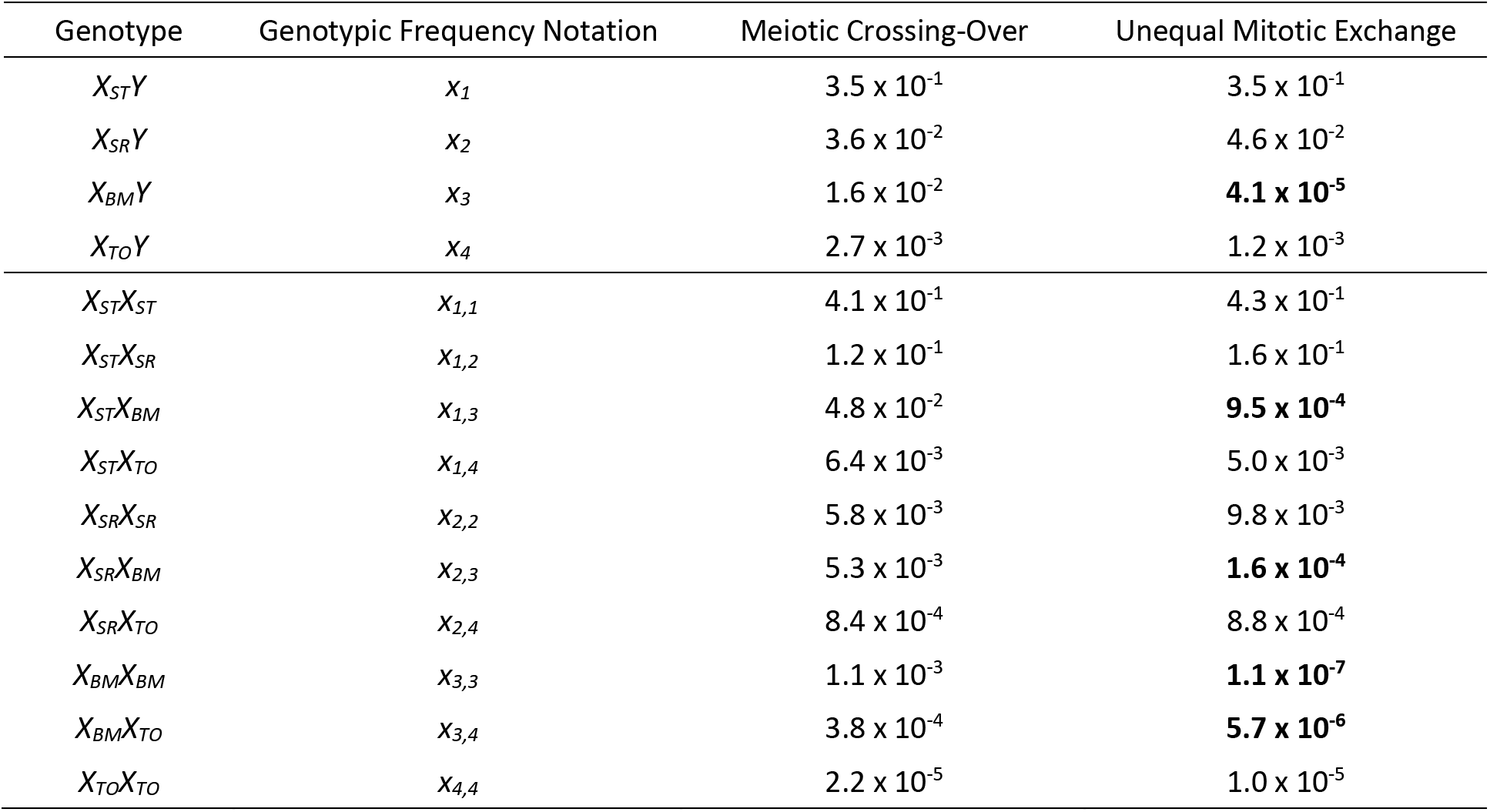
Numerical solutions for equilibrium genotypic frequencies in *D. pseudoobscura Sex Ratio* system. The equilibrium frequencies are listed for all genotypes (4 male hemizygous states, 4 female homozygous states, and 6 female heterozygous states) in the recombination extensions of Edwards (1961) model of *SR* evolution. Full system of 14 equations is derived in the Appendix by assuming recombination was either reciprocal-exchange caused by meiotic crossing-over in the primary oocyte, or alternately caused by unequal mitotic exchange in female germline stem cells. Under alternate assumptions, the largest differences all involve the genotypes with *X_BM_* and are highlighted in bold.

Assuming *X_SR_/X_ST_* recombination was due to meiotic crossing-over, the modeled equilibrium frequencies of recombinant *X* chromosomes were 0.045 (*X_BM_*) and 0.0065 (*X_TO_*), with *SR* inversion linkage disequilibrium *r*^2^ = 0.62 (Figure 4A). These modeling results can be evaluated by comparing to the gold standard of Beckenbach’s (1996) direct cytological observation of *SR* inversion complement for 694 *X* chromosomes sampled from Tucson and Bear Creek, Arizona, USA. The *SR* population genetic model assuming meiotic recombination compared to direct natural population observations reveals: 1) incorrect prediction of the rank order of recombinant *X* chromosomes, 2) inaccurate estimates of the natural population *X_SR_*, *X_BM_*, and *X_TO_* frequencies, and 3) a large discrepancy in linkage disequilibrium of *SR* inversions (Table 7). Although Fuller *et al.* 2020 used a different modeling framework, it was the discrepancy in linkage disequilibrium of *SR* inversions that precipitated our previously published conclusion of strong epistatic selection acting to maintain the full 26 Mb *SR* haplotype.

**Figure 4.**
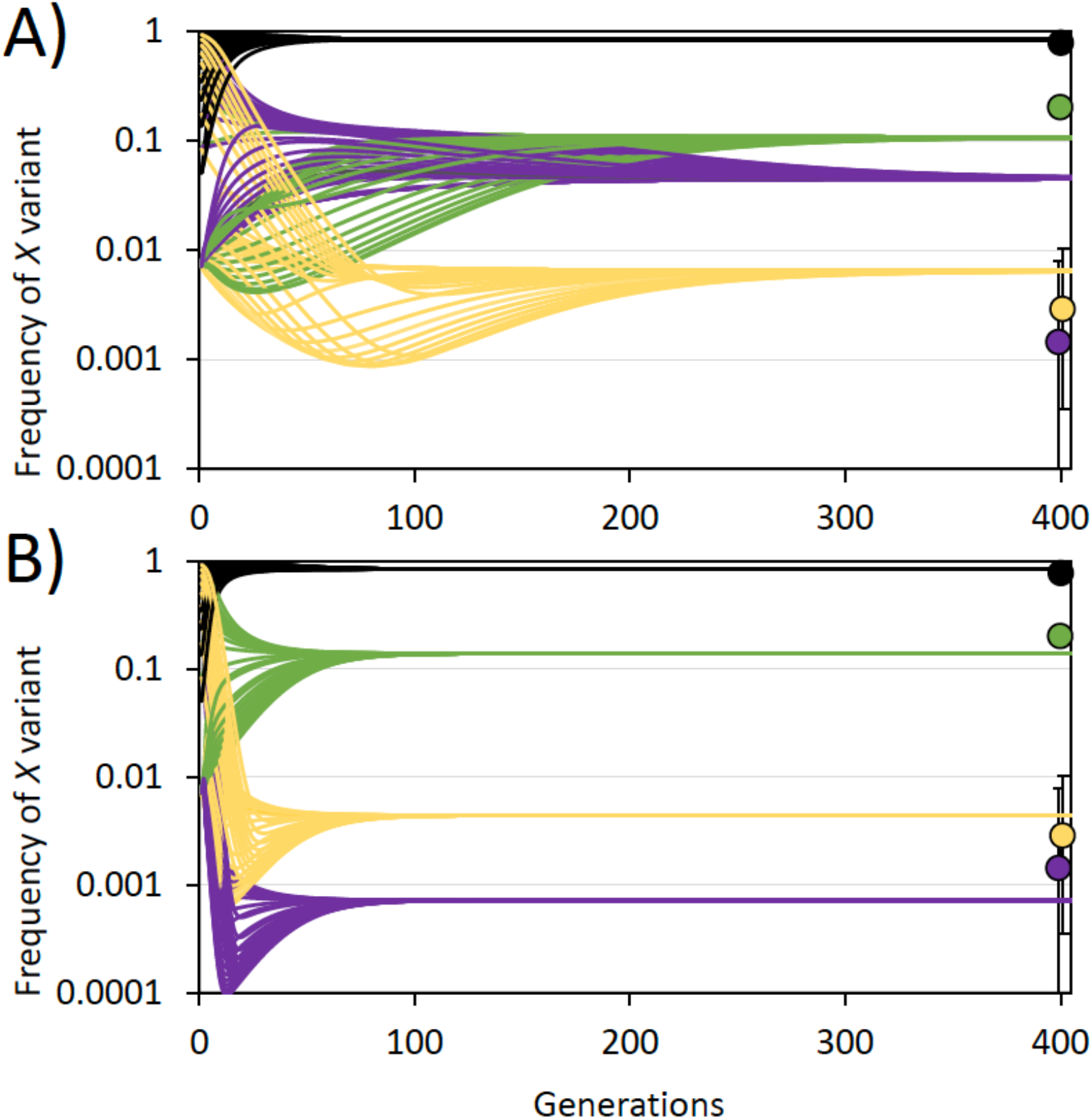
Modeled evolutionary trajectories for *X* chromosome variants in *D. pseudoobscura*. Change in population frequency of *X_ST_* (black), *X_SR_* (green), *X_BM_* (purple), and *X_TO_* (orange) chromosomes with initial frequencies ranging from 0.01 to 0.99. For comparison, Beckenbach’s (1996) observations from Arizona, USA are given as color coded points on the far right with exact binomial 95% confidence intervals. A) Extension of Edwards (1961) population genetic model assuming recombination is reciprocal-exchange caused meiotic crossing-over. B) The same model incorporating experimentally determined asymmetries in recombination due to unequal mitotic exchange. The Edwards (1961) model incorporating unequal mitotic exchange is a much better fit to the frequencies observed in the wild (see Table 7).

To model *X_SR_/X_ST_* recombination as unequal mitotic exchange, independent complementary recombinant class-specific rates (*c*_3_ and *c*_4_) were calculated from experimental data reported in the present study using only the protected heterozygous state (*i.e.*, from *F_2_* female progeny only in Figure 3C) (*c*_3_ = 0.0072, *c*_4_ = 0.018). Similarly, hemizygous relative viabilities for the complementary recombinants were calculated as the experimentally observed proportional deficit of recombinants observed in *F_2_* male versus *F_2_* female progeny in Figure 3C (*v*_3_ = 0.05, *v*_4_ = 0.39). Solving the system of equations (Equation A2 in Appendix) with new parameter values reflecting asymmetries in recombination due to unequal mitotic exchange revealed the largest changes in equilibrium frequencies all involve genotypes containing *X_BM_* (bold values Table 6 column “Unequal Mitotic Exchange”).

Assuming the experimentally observed *X_SR_/X_ST_* recombination was due to unequal mitotic exchange, the modified model described above produced equilibrium frequencies of 0.00072 (*X_BM_*) and 0.0044 (*X_TO_*), both of which are within the 95% confidence intervals of Beckenbach’s (1996) sampled frequencies (Figure 4B). Further benchmarking against Beckenbach (1996) demonstrated the *SR* population genetic model incorporating experimentally determined recombination asymmetries could: 1) correctly predict rank order of recombinant *X* chromosome frequencies, 2) improve fit to natural population *X_SR_*, *X_BM_*, and *X_TO_* frequencies, and 3) closely match *SR* inversion linkage disequilibrium in the wild, thereby eliminating the need to invoke any additional selection (Table 7).

**Table 7.** Observed and modeled *D. pseudoobscura X* chromosome variant frequencies and linkage disequilibrium of *SR* haplotype inversions. Direct cytological observations of Beckenbach (1996) were compared to the modeled frequencies and linkage disequilibrium from Fuller *et al.* (2020) as well as the recombination extensions of Edwards (1961) derived in the Appendix. The best fit to natural population frequencies of *X_ST_* and *X_SR_* chromosomes, their recombinants *X_BM_* and *X_TO_*, and linkage disequilibrium (*r*^2^) of *SR* haplotype inversions was the model for *Sex Ratio* evolution incorporating unequal mitotic exchange through experimentally determined asymmetries in recombination.

|  | Beckenbach (1996)<br>Direct Cytological<br>Observations | Fuller *et al.* (2020)<br>Decay Model for Gametic<br>Phase Disequilibrium | Edwards (1961)<br>SR Model Assuming<br>Meiotic Crossover | Edwards (1961)<br>SR Model Assuming<br>Unequal Mitotic Exchange |
| --- | --- | --- | --- | --- |
| $X_{ST}$ | 0.79 | 0.64 | 0.84 | 0.85 |
| $X_{SR}$ | 0.20 | 0.04 | 0.11 | 0.14 |
| $X_{BM}$ | 0.001 | 0.16 | 0.045 | 0.001 |
| $X_{TO}$ | 0.003 | 0.16 | 0.007 | 0.004 |
| $r^2$ | 0.97 | 0.00 | 0.62 | 0.97 |

Whereas the statistical analysis of the experiment strongly rejects the null hypothesis that meiotic crossing-over generated the recombination dataset, it is the modeling of natural population frequencies in the preceding paragraphs that reveals adopting the alternative hypothesis of unequal mitotic exchange is important for understanding the evolutionary dynamics of the *D. pseudoobscura SR* system. Of the three main experimental results, 1) higher recombination rates, 2) clustered recombination events, and 3) asymmetric recovery of complementary recombinant classes, it appears that the last of these has the greatest importance in explaining the equilibrium frequencies of *X* chromosome variants in natural populations. This analysis suggests incorporating the experimentally determined deficit of hemizygous recombinants into the population model in the form of recombination-induce viability defects is sufficient to explain *X_BM_* and *X_TO_* rarity in the wild, thereby eliminating the need to invoke additional epistatic selection to maintain the highly differentiated 26 Mb *SR* haplotype.

### Conclusion

The recombination rate in the 6.6 Mb colinear region of *D. pseudoobscura X_SR_/X_ST_* heterozygotes was revised upwards to 0.00779 (92 recombinants in 11,808 progeny) using an improved experimental design. Statistical analysis of rate heterogeneity revealed that the majority of these recombination events did not occur as independent meiotic crossover event in primary oocytes, but rather they were clustered in a manner consistent with unequal mitotic exchange in the female GSCs. Incorporating the experimentally observed asymmetries in recombination into a sex-specific, genotypic models of *SR* evolution provided a substantially better fit to natural population frequencies and eliminates the need to invoke any additional selection to maintain the 26 Mb *D. pseudoobscura SR* haplotype in the wild.

## Data Availability

*D. pseudoobscura* stocks listed in Table 1 and further described in Reagents Table in File S1 are available upon request..The author affirms that all data necessary for confirming the conclusions of the article are present within the article, figures, and tables. Raw counts from the recombination experiments are provided as a supplement in File S3.

## Acknowledgements

Data analysis and manuscript preparation was performed at Stowers Institute for Medical Research supported by NRSA fellowship NIH F32 GM134707 awarded to Spencer Koury. The experiment was performed at University of Utah with funds provided from NIH 1R01 GM115914 awarded to Nitin Phadnis. Research funders and sponsor had no role in design, execution, analysis, interpretation, and reporting of this study. The manuscript benefitted from suggestions by Nitin Phadnis, Sarah Gross, and Shelley Reich of University of Utah.

## APPENDIX

### Population Genetic Modeling

To determine the implications of the experimental results for natural populations, the Edwards (1961) model of *Sex Ratio* (*SR*) in *Drosophila pseudoobscura* was extended to include recombination. Following the original model, female genotypes for standard arrangement *X* chromosomes and the multiply-inverted *SR* chromosome variants are *X_ST_/X_ST_, X_ST_/X_SR_*, and *X_SR_/X_SR_* with frequencies denoted by *x*_1,1_, *x*_1,2_, and *x*_2,2_, respectively. The male genotypes are *X_ST_/Y* and *X_SR_/Y* with respective frequencies *x*_1_ and *x*_2_, such that *x*_2_ + *x*_2_ + *x*_1,1_ + *x*_1,2_ + *x*_2,2_ = 1. Let the relative viabilities of these five genotypes be *v*_1_, *v*_2_, *v*_1,1_, *v*_1,2_, *v*_2,2_, with relative fertilities *f*_1_, *f*_2_, *f*_1,1_, *f*_1,2_, *f*_2,2_, where the subscript denotes the corresponding genotype. To model *X*-linked segregation distortion, male genotypes produce a proportion of female offspring *k_i_* with *k*_1_ = 0.5 representing mendelian transmission of *X_ST_* chromosomes and *k*_2_ = 1.0 representing complete distortion of *X_SR_* chromosomes.

To incorporate recombination into the Edwards (1961) model, let exchange in the 6.6 Mb colinear region located between medial and terminal inversions of *SR* chromosomes occur with frequency *c*_1,2_ in the double heterozygotes *X_SR_/X_ST_*. Exchange in the colinear region generates two new recombinant *X* chromosomes: *X_BM_*, with basal and medial inversions and approximately 2/3 of the 26 Mb *Sex Ratio* haplotype, and *X_TO_*, with carries the terminal inversion only and about 1/3 of the 26 Mb *Sex Ratio* haplotype. Male genotypic frequency of *X_BM_/Y* and *X_TO_/Y* are denoted *x*_3_ and *x*_4_, respectively, with corresponding relative viabilities and relative fertilities of *v*_3_, *v*_4_ and *f*_3_, *f*_4_. Following this notation, the female genotypes *X_ST_/X_BM_, X_ST_/X_TO_, X_SR_/X_BM_, X_SR_/X_TO_, X_BM_/X_BM_, X_BM_/X_TO_*, and *X_TO_/X_TO_* have frequencies *x*_1,3_, *x*_1,4_, *x*_2,3_, *x*_3,3_, *x*_3,4_, *x*_4,4_ and corresponding relative viabilities *v*_1,3_, *v*_1,4_, *v*_2,3_, *v*_3,3_, *v*_3,4_, *v*_4,4_ and relative fertilities *f*_1,3_, *f*_1,4_, *f*_2,3_, *f*_3,3_, *f*_3,4_, *f*_4,4_. All genotypes, genotypic frequencies, and their corresponding parameters (relative viability, relative fertility, male germline *X* chromosome transmission, and female germline *X* chromosome recombination) are given in the mating table provided in File S4.

To calculate genotypic frequencies in the next generation, a mating table was created that lists all genotypic frequencies of offspring produced from all 40 mating types (4 possible male genotypes × 10 possible female genotypes). The full table (40 parental mating types generating 14 offspring genotypes) is given in supplemental File S4. After correcting for the frequency and fertility of each parental mating type, as well as relative viability of each offspring class, the genotypic frequencies of reproductively mature adults in the next generation are given by system of equations:

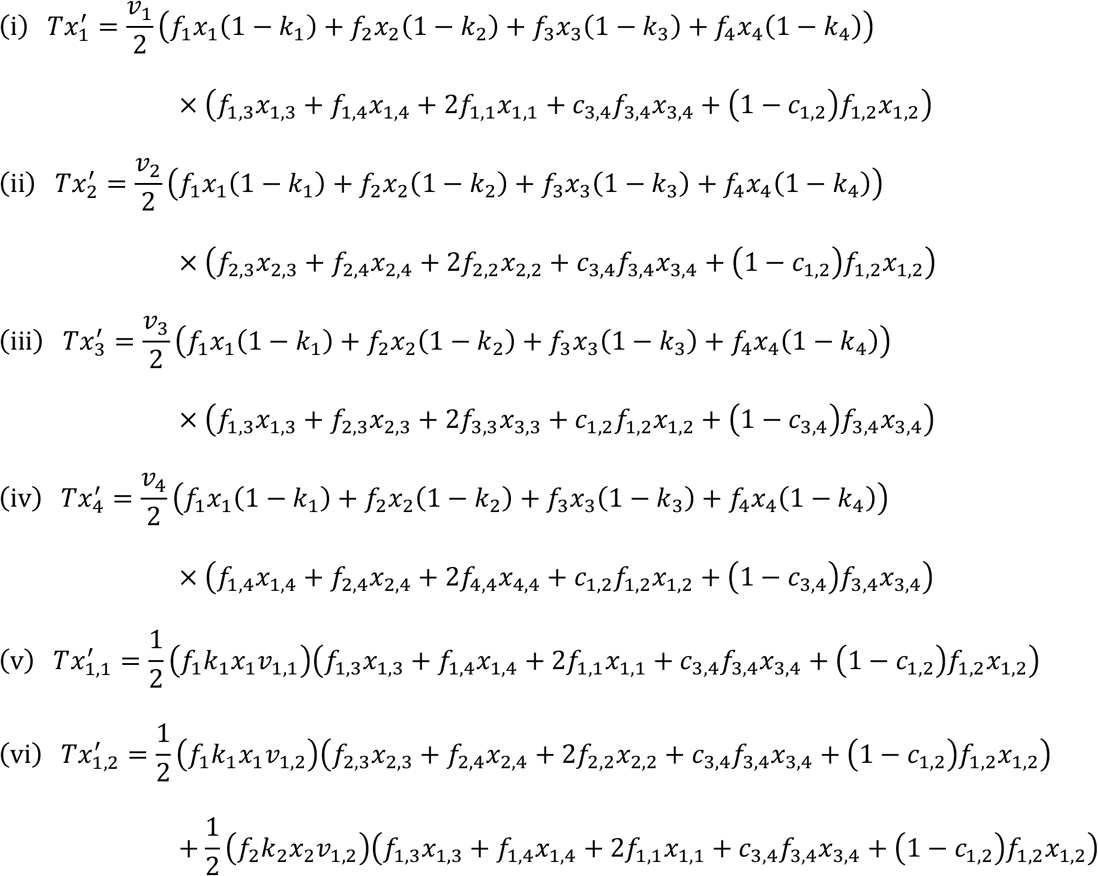

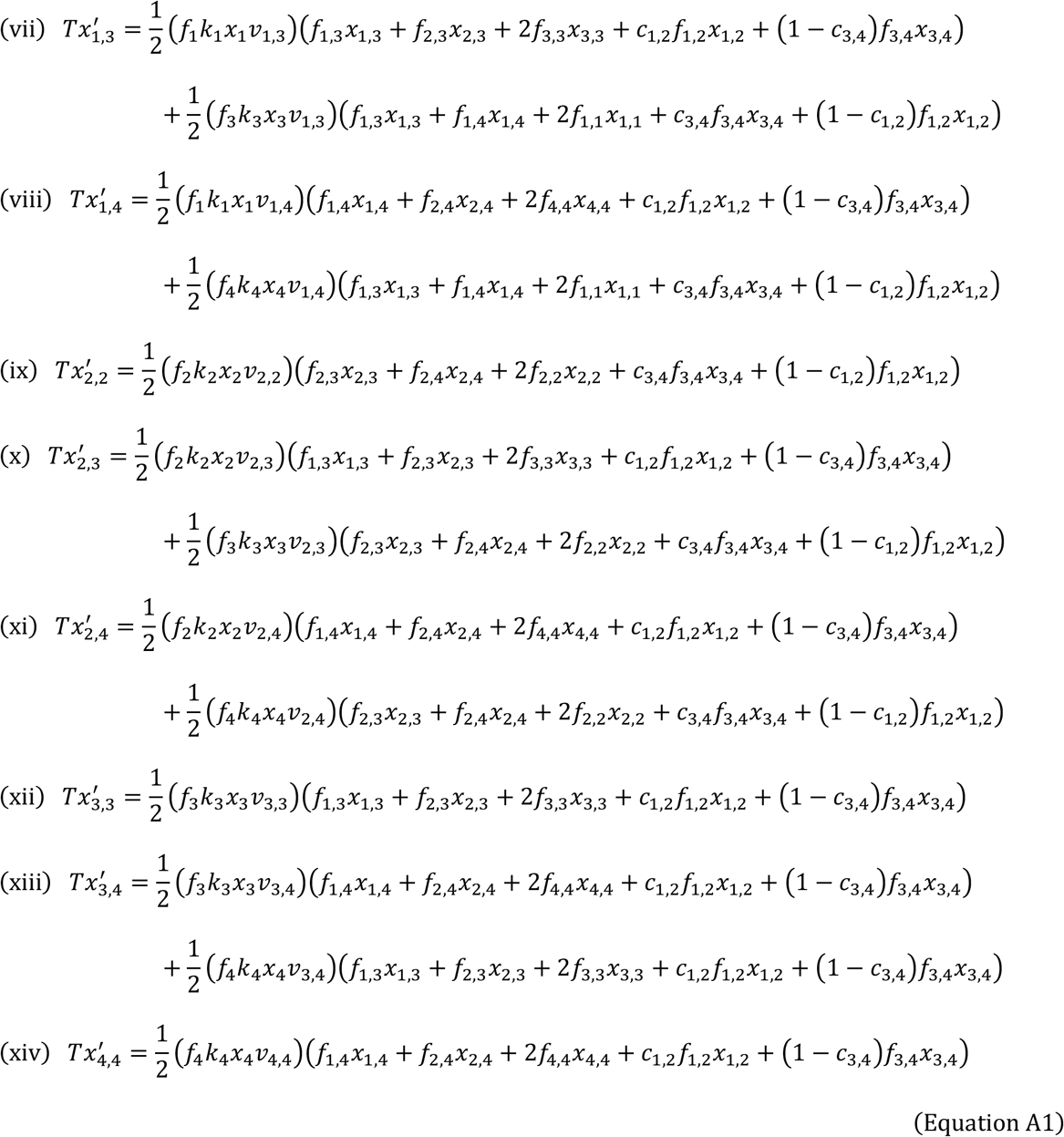

where *T* is the sum of the right side of the system of equations and the next generation’s genotypic frequencies are denoted with primes. Iteration of this single generation system allows graphical analysis of short-term evolution of *D. pseudoobscura Sex Ratio* system.

The behavior of this model was examined with experimentally determined parameters and several simplifying assumptions. First, relative viability and fertility for genotypes involving *X_ST_* and *X_SR_* were taken from direct measurements in Curtsinger and Feldman (1980) and are assumed to be recessive such that *v*_1,2_ = *f*_1,2_ = 1. Second, relative viabilities and fertilities were products of multiplicative fitness effects; such that when the 26 Mb *SR* haplotype was disrupted by recombination, then *v*_2,2_ = *v*_2,3_ *v*_2,4_ and *f*_2,2_ = *f*_2,3_ *f*_2,4_. Furthermore, relative viability and fertility of recombinant *X* chromosomes in the homozygous state was proportional to the fraction of the 26 Mb *SR* haplotype retained, such that 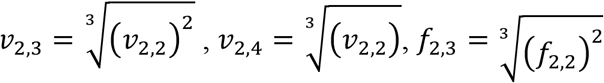, and 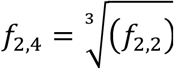. Following the same logic for hemizygote relative viability and fertilities were 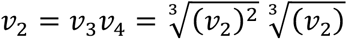 and 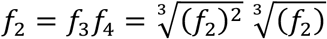. All full list numerical values for viability and fertility parameters are given in Table A1. Third, the *X*-linked segregation distortion (male genotype-specific proportion of female offspring sired) were *k*_1_ = 0.5, *k*_2_ = 0.5, *k*_3_ = 0.8, and *k*_4_ = 1.0 based on experimental estimates. Here, an averaged *k*_3_ was used to account for polymorphic autosomal suppressors in natural populations that can alter the distortion strength of *X_BM_* (unpublished results). Finally, recombination rates were based on the experimental results reported in this study, however, two different models of recombination (reciprocal exchange due to meiotic crossing over and unequal mitotic exchange in female germline stem cells) were investigated.

Meiotic crossing-over was modeled as reciprocal-exchange using the experiment-wide rate of 0.00779 events per offspring for both directions of recombination *c*_1,2_ (producing *X_BM_* and *X_TO_*) and *c*_3,4_ (producing *X_ST_* and *X_SR_*). The “back” recombination rate *c*_3,4_ (heterozygous *X_BM_/X_TO_* females generating *X_ST_* or *X_SR_* progeny) has not been experimentally estimated. However, the relevant female genotype *x*_3,4_ only occurs on the order of 1 in 1,000,000 individuals in the natural population. Meiotic crossing-over produces system of equations A1 derived above.

Unequal mitotic exchange in the female germline stem cells was modeled by substituting the “forward” recombination rate (*c*_1,2_ producing *X_BM_* and *X_TO_* from *X_ST_/X_SR_* female heterozygotes) with complementary class-specific rates *c*_3_ (generating only *X_BM_*) and *c*_4_ (generating only *X_TO_*) into the relevant columns of the mating table. Complementary class-specific rates (*c*_3_ = 0.00715 and *c*_4_ = 0.0182) were calculated from experimental recovery rates in the protected heterozygous state (*i.e.*, from female progeny only). The “back” recombination rate *c*_3,4_ was not altered as the relevant genotype *c*_3,4_ was exceedingly rare. Finally, relative viability of hemizygous *X_BM_* and *X_TO_* were estimated by deficit of recombinant male versus female progeny observed in experimental data (*v*_3_ = 0.050 and *v*_4_ = 0.392). Therefore, the modified system of equations for unequal mitotic exchange were:

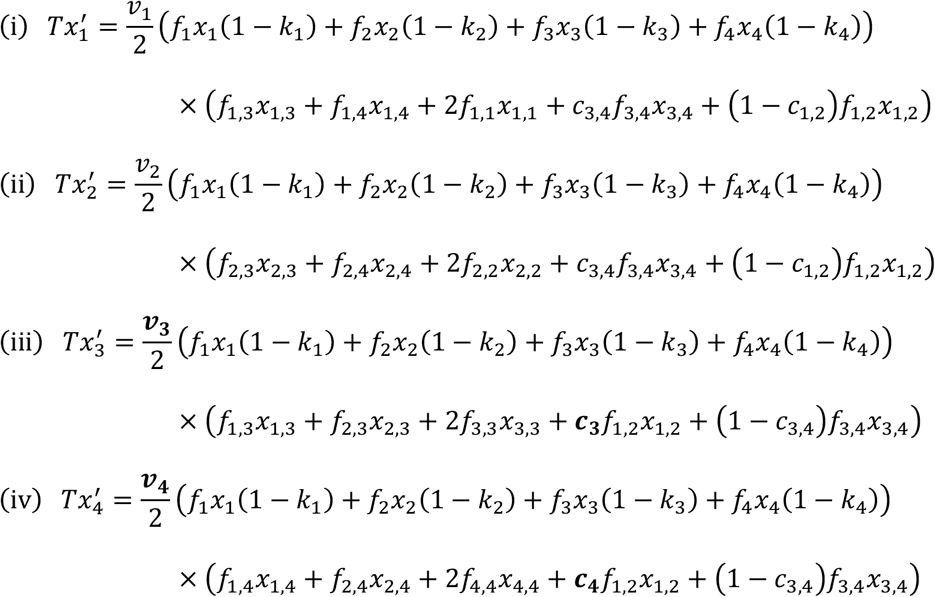

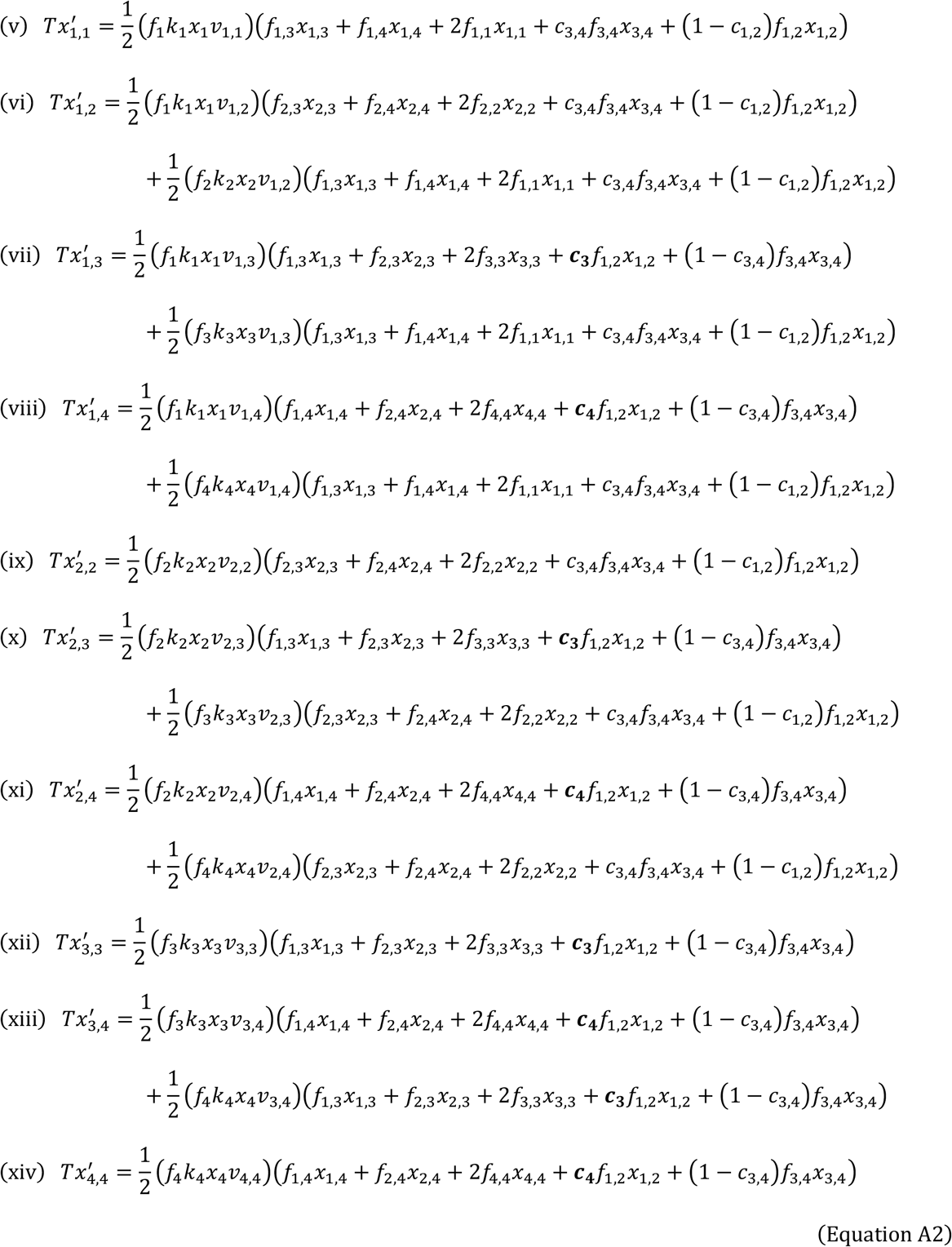

where, as before, *T* is the sum of the right side of equations, primes denote next generation’s genotypic frequencies, and the altered terms (*c*_3_, *c*_4_, *v*_3_, and *v*_4_) are bolded for convenience.

Thorough equilibrium and stability analysis of the full 14 variable population genetic model is beyond the scope of this paper. Instead, equilibrium frequencies were solved for by iteration of the system of recursion equations. Initial genotypic frequencies were taken from Beckenbach’s (1996) direct cytological observation of *SR* inversion complement for 694 *X* chromosomes sampled from Tucson and Bear Creek, Arizona, USA. The equilibrium frequencies are reported in main text Table 6. This procedure was then repeated with a range of initial allele frequencies (0.01-0.99) for *X_SR_*, *X_BM_*, and *X_TO_* under both alternative scenarios: meiotic crossing-over in oocyte or unequal mitotic exchange in germline stem cells. In all cases, the genotypic frequencies converged on the same values in less than 1000 generations (main text Figure 4).

## APPENDIX TABLE

**Table A1.** Viability and fertility parameter values for the recombination extension of Edwards (1961) model. Parameter are classified as “defined” if their value is set by the modeling framework, “empirical” if directly measured in Curtsinger and Feldman (1980), and “calculated” if based on proportional division of Curtsinger and Feldman (1980) values.

| Genotype | Viability |  |  | Fertility |  |  |
| --- | --- | --- | --- | --- | --- | --- |
|  | Parameter | Estimate | Method | Parameter | Estimate | Method |
| $X_{ST}Y$ | $v_1$ | 1.02 | Empirical | $f_1$ | 1.00 | Empirical |
| $X_{SR}Y$ | $v_2$ | 0.71 | Empirical | $f_2$ | 1.03 | Empirical |
| $X_{BM}Y$ | $v_3$ | 0.80 | Calculated | $f_3$ | 1.02 | Calculated |
| $X_{TO}Y$ | $v_4$ | 0.89 | Calculated | $f_4$ | 1.01 | Calculated |
| $X_{ST}X_{ST}$ | $v_{1,1}$ | 1.24 | Empirical | $f_{1,1}$ | 0.87 | Empirical |
| $X_{ST}X_{SR}$ | $v_{1,2}$ | 1.00 | Defined | $f_{1,2}$ | 1.00 | Defined |
| $X_{ST}X_{BM}$ | $v_{1,3}$ | 1.07 | Calculated | $f_{1,3}$ | 0.95 | Calculated |
| $X_{ST}X_{TO}$ | $v_{1,4}$ | 1.15 | Calculated | $f_{1,4}$ | 0.91 | Calculated |
| $X_{SR}X_{SR}$ | $v_{2,2}$ | 0.55 | Empirical | $f_{2,2}$ | 0.68 | Empirical |
| $X_{SR}X_{BM}$ | $v_{2,3}$ | 0.67 | Calculated | $f_{2,3}$ | 0.77 | Calculated |
| $X_{SR}X_{TO}$ | $v_{2,4}$ | 0.82 | Calculated | $f_{2,4}$ | 0.88 | Calculated |
| $X_{BM}X_{BM}$ | $v_{3,3}$ | 0.72 | Calculated | $f_{3,3}$ | 0.73 | Calculated |
| $X_{BM}X_{TO}$ | $v_{3,4}$ | 1.00 | Calculated | $f_{3,4}$ | 1.00 | Calculated |
| $X_{TO}X_{TO}$ | $v_{4,4}$ | 0.95 | Calculated | $f_{4,4}$ | 0.80 | Calculated |

